## Supplementary Information for "Cell crowding activates pro-invasive mechanotransduction pathway in high-grade DCIS via TRPV4 inhibition and cell volume reduction"

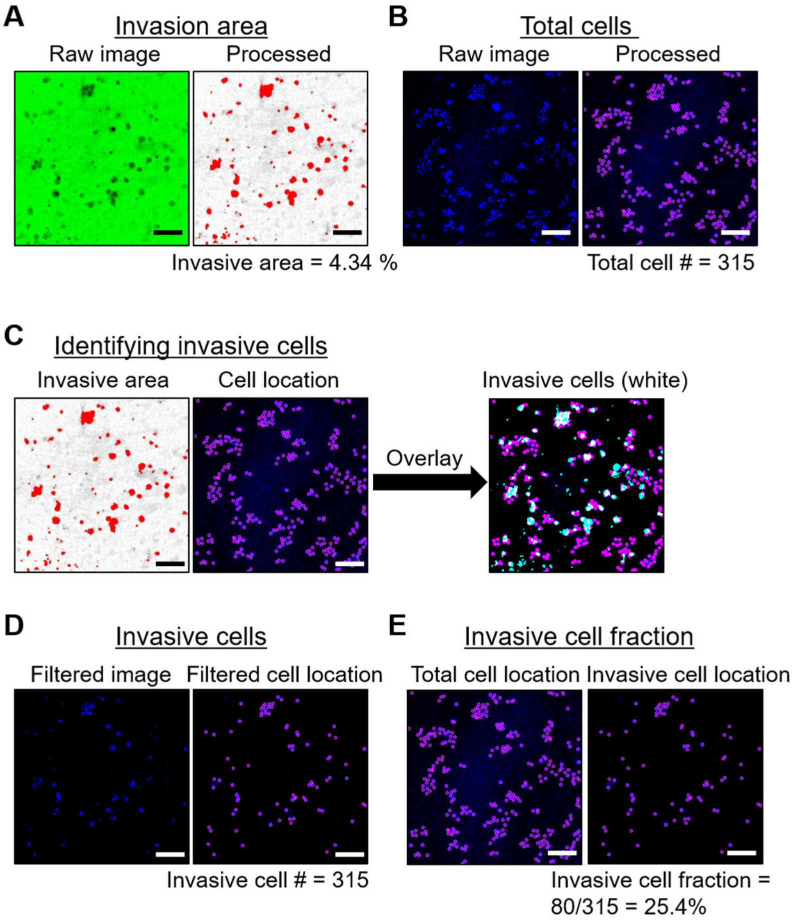


**Supplementary Figure 1. Quantifying the Invasive cell fraction using a 2D polyacrylamide hydrogel-based invasion assay. (A)** Green fluorescent gelatin images were thresholded to highlight invasion areas (in red) due to degradation (visible as dark regions in the original image). Here, 4.34% of the total area indicated cell invasion. **(B)** DAPI images (in blue, showing nuclei locations) were processed with the Trackmate plugin in Image J to detect individual DAPI spots (displayed in purple). The total cell count in this instance was estimated at 315. **(C)** The highlighted invasive areas from **(A)** served as masks for the DAPI locations, selecting only the invasive cells (those in the white regions resulting from the overlay between cell positions in purple and invasive zones in cyan). **(D)** The resultant images display the locations of invasive cells in purple. The invasive cell count from this image was 80. **(E)** The fraction of invasive cells was determined by comparing the number of invasive cells (from **D**) with the overall cell count (from **B**). Thus, the invasive cell fraction was 25.4% (80 out of 315). Scale bar = 100 μm.

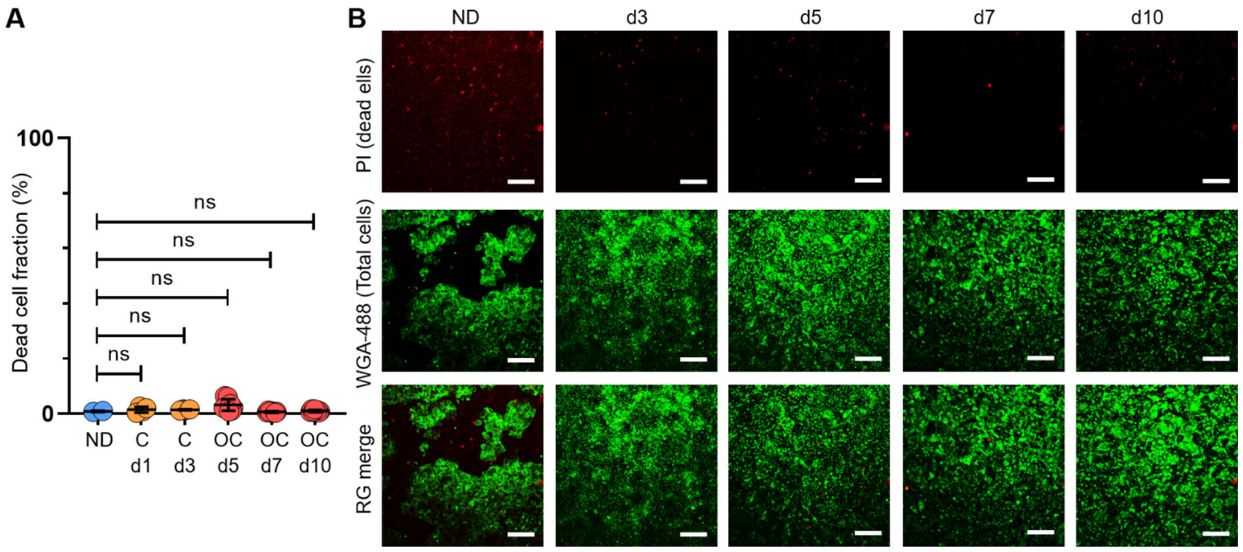


**Supplementary Figure 2. MCF10DCIS.com cells exhibited comparable viability under OC conditions to those under ND conditions. (A-B)** Live cells seeded at ND (~70%) (n=2) were cultured for 1 (n=4), 3 (n=4), 5 (n=8), 7 (n=8), and 10 (n=8) days post-confluence (denoted as "d"). Cells were stained with propidium iodide (PI, red) to mark dead cells, followed by WGA-488 (green) to label nuclei. After staining, cells were fixed and imaged using confocal microscopy (4X), with red and green signals quantified via the TrackMate plugin in ImageJ. The percentage of dead cells, based on PI-positive signals, is presented in **A**. Both confluent (day 3) and OC (day 5–10) MCF10DCIS.com cells showed similar viability to ND cells, with less than 1% cell death (0.85 ± 0.25%). On average, the fraction of dead cells was 1.58 ± 0.98%, confirming that cell crowding does not additionally induce cell death. Representative PI/WGA-488/merged images from days 1, 3, 5, 7, and 10 are shown in **B.** Scale bar = 100 μm. For the t-test, we employed a nonparametric approach using the Mann-Whitney test with a two-tailed p-value, which was used throughout the manuscript. The statistical significance levels are denoted as follows: ****: $p < 0.0001$, ***: $p < 0.001$, **: $p < 0.01$, *: $p < 0.05$, ns: $p > 0.05$.

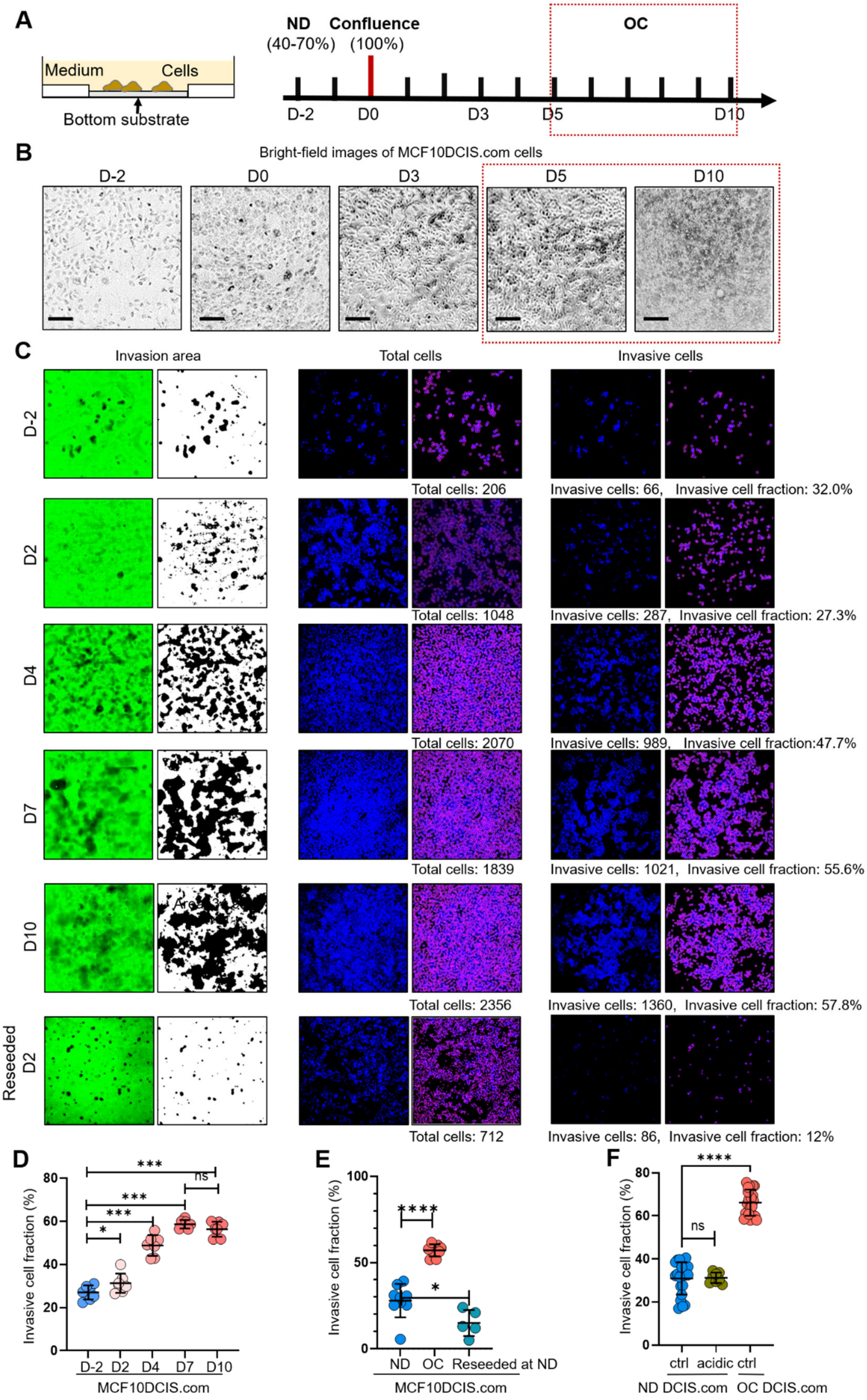
A
Medium
Cells
Bottom substrate
ND
(40-70%)
Confluence
(100%)
OC
D-2
D0
D3
D5
D10
B
Bright-field images of MCF10DCIS.com cells
D-2
D0
D3
D5
D10
C
Invasion area
Total cells
Invasive cells
D-2
Total cells: 206
Invasive cells: 66, Invasive cell fraction: 32.0%
D2
Total cells: 1048
Invasive cells: 287, Invasive cell fraction: 27.3%
D4
Total cells: 2070
Invasive cells: 989, Invasive cell fraction:47.7%
D7
Total cells: 1839
Invasive cells: 1021, Invasive cell fraction: 55.6%
D10
Total cells: 2356
Invasive cells: 1360, Invasive cell fraction: 57.8%
Reseeded D2
Total cells: 712
Invasive cells: 86, Invasive cell fraction: 12%
D
Invasive cell fraction (%)
D-2
D2
D4
D7
D10
MCF10DCIS.com
E
Invasive cell fraction (%)
ND
OC
Reseeded at ND
MCF10DCIS.com
F
Invasive cell fraction (%)
ctrl
acidic
ctrl
ND DCIS.com
OC DCIS.com

**Supplementary Figure 3. Time window for cell crowding conditions in vitro. (A)** MCF10DCIS.com cells were grown to: normal density (ND; 40%–70%), confluence (100%), and overconfluence (OC). The time points for these growth stages were two days before confluence (D-2) for ND, day 0 (D0) for confluence, and days 5–10 after confluence (D5) for OC. We selected the cell crowding condition (OC conditions) during days 5–10 as cell morphology (**B**) and invasiveness (**C**) reached equilibrium after day 5. **(B)** Brightfield microscopy images of MCF10DCIS.com cells on the indicated days. Cells exhibited significant compaction starting from day 5. Scale bar = 100 μm. (**C-D**) Time-dependent invasiveness of MCF10DCIS.com cells as they progressed to OC. The cell invasiveness increased until day 5 and plateaued between days 5 and 10. When these cells were reseeded at normal density on day 2 (reseeded D2), their invasiveness decreased to levels similar to the original ND cells. **(E)** Formerly OC cells reseeded at ND showed a slight reduction in the fraction of invasive cells (light teal circles) to approximately 15%, comparable to ND cells (blue circles) before exposure to OC conditions. These results suggest that the OC-induced increase in invasiveness is largely reversible. **(F)** To test if acidity of OC cell media, despite frequent changing, contributed to increased invasiveness of DCIS.com cells, we used acidic OC media (day 7) to treat ND DCIS.com cells for two days. Conditioned media did not alter invasiveness of ND DCIS.com cells. For the t-test, we employed a nonparametric approach using the Mann-Whitney test with a two-tailed p-value, which was used throughout the manuscript. The statistical significance levels are denoted as follows: ****: $p < 0.0001$, ***: $p < 0.001$, **: $p < 0.01$, *: $p < 0.05$, ns: $p > 0.05$.

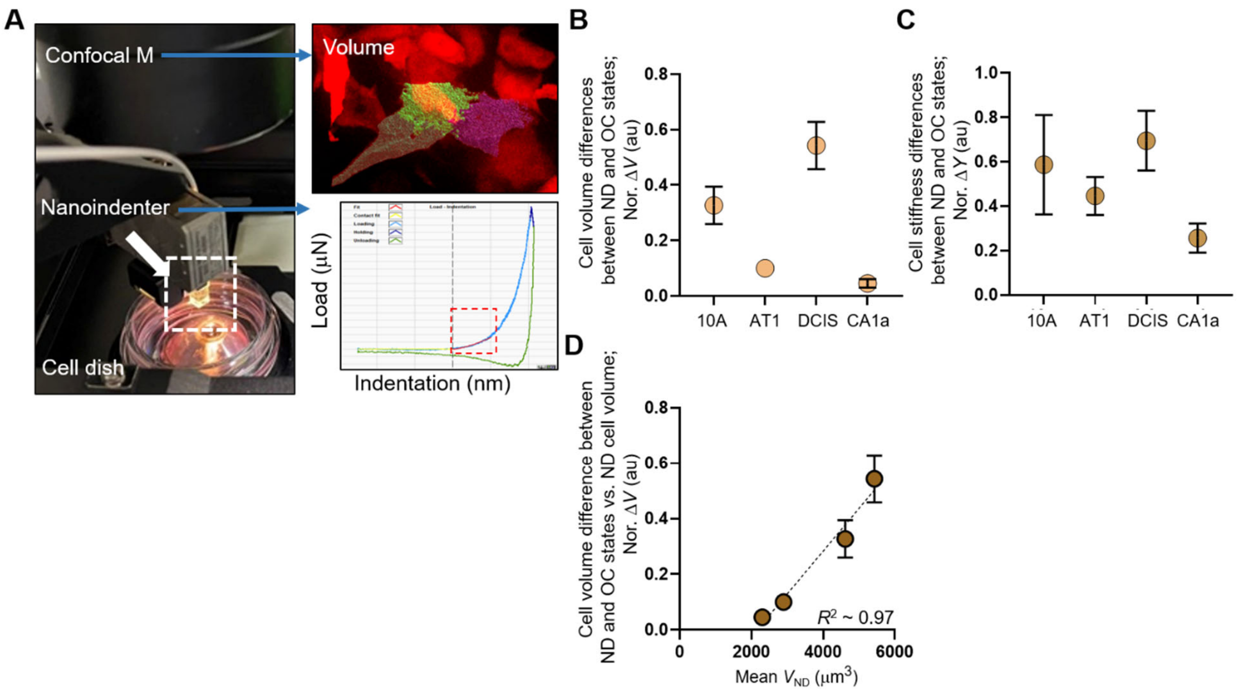


**Supplementary Figure 4. Increased invasiveness of MCF10DCIS.com cells correlates with cell volume plasticity, not the acidity of cell media. (A)** Cell crowding induced cell volume and stiffness changes assessed by our confocal microscope, which captures 3D volume of single RFP-expressing cells (right image) and includes a nanoindenter device (Chiaro, Optics11life) that can indent a single cell (load vs indentation curve) to extract Young's modulus in the elastic regime (red dashed box) using a Hertzian model. The indentation probe has a spring constant and tip diameter of ~0.24 N/m and 10 μm, respectively. We confirmed that RFP expression did not alter cell volume. **(B-C)** Cell volume and stiffness differences between ND and OC conditions calculated using the cell volume and stiffness data in Fig. 2A and 2C were normalized to the ND cell volume **(B)** or stiffness **(C)**, with the highest changes observed in high-grade MCF10DCIS.com cells. **(D)** Normalized volume change (Nor. Δ$V$) linearly scaled with ND cell volume (mean $V_{ND}$), with an $R^2$ value of approximately 0.97, signifying a highly linear relationship. ****: $p < 0.0001$, ***: $p < 0.001$, **: $p < 0.01$, *: $p < 0.05$, ns: $p > 0.05$.

| **Gene names MCF10DCIS.com cells** | **Protein Name (MCF10DCIS.com)** | **OC/ND ratio (MCF10DCIS.com)** |
|---|---|---|
| **TRPV4** | Transient Receptor Potential Vanilloid 4 | 153.01 |
| IVL | Involucrin | 55.35 |
| **SCN11A;SCN5A;SCN10A;SCN9A** | Sodium Voltage-Gated Channel Alpha Subunit 11; Sodium Voltage-Gated Channel Alpha Subunit 5; Sodium Voltage-Gated Channel Alpha Subunit 10; Sodium Voltage-Gated Channel Alpha Subunit 9 | 41.82 |
| HNRNPM | Heterogeneous Nuclear Ribonucleoprotein M | 26.91 |
| COPG2 | Coatomer Protein Complex Subunit Gamma 2 | 18.33 |
| S100P | S100 calcium-binding protein P | 18.01 |
| MRPL16 | Mitochondrial Ribosomal Protein L16 | 14.43 |
| IDH2 | Isocitrate Dehydrogenase [NADP] Mitochondrial | 8.68 |
| EML4 | Echinoderm Microtubule-Associated Protein-Like 4 | 7.94 |
| GSN | Gelsolin | 7.89 |
| CKAP4 | Cytoskeleton-Associated Protein 4 | 7.82 |
| PRSS8 | Serine Protease 8 | 6.93 |
| MFI2 | Melanotransferrin | 6.85 |
| CEACAM1 | Carcinoembryonic Antigen-Related Cell Adhesion Molecule 1 | 6.84 |
| ADGRL3;LPHN3 | Adhesion G protein-coupled receptor L3; Latrophilin-3 | 6.82 |
| HNRNPC | Heterogeneous Nuclear Ribonucleoprotein C | 6.78 |
| RPL7L1 | Ribosomal Protein L7-Like 1 | 6.54 |

| | | |
|---|---|---|
| NDUFV1 | NADH:Ubiquinone Oxidoreductase Core Subunit V1 | 6.19 |
| HIST1H2A;HIST1H2AH;HIST1H2AF;HIST1H2AJ;HIST1H2AG;HIST1H2AA3;HIST2H2AA3;H2AFX | Histone H2A family | 6.07 |
| GTF2I | General Transcription Factor II-I | 5.88 |
| ERP44 | Endoplasmic Reticulum Protein 44 | 5.81 |
| **KCNN4** | Potassium Intermediate Conductance Calcium-Activated Channel 4 | 5.61 |
| NDUFB9 | NADH:Ubiquinone Oxidoreductase Subunit B9 | 5.59 |
| RPL32 | Ribosomal Protein L32 | 5.47 |

**Supplementary Table 1.** Gene and protein names that showed more than a 5-fold increase in plasma membrane association under OC conditions relative to ND conditions in MCF10DCIS.com cells were identified by mass spectrometry. Ion channels among these are highlighted in bold.

| Gene MCF10A cells | Protein Name (MCF10A) | OC/ND ratio (MCF10A) | Gene MCF10AT1 cells | Protein Name (MCF10AT1) | OC/ND ratio (MCF10AT1) | Gene MCF10CA1a cells | Protein Name (MCF10CA1a) | OC/ND ratio (MCF10CA1a) |
|---|---|---|---|---|---|---|---|---|
| SCEL | Scellicin | 5318.64 | HTRA2 | High temperature requirement A2 | 7.36 | ACAA2 | Acetyl-CoA acyltransferase 2 | 82.61 |
| ALDH7A1 | Aldehyde dehydrogenase 7 family member A1 | 3111.33 | PMPCB | Peptidase (mitochondrial processing) beta | 5.86 | **ATP2B4** | Plasma membrane calcium-transporting ATPase 4 | 37.51 |
| ECH1 | Enoyl CoA hydratase 1 | 2711.84 | CLPP | ATP-dependent Clp protease proteolytic subunit | 5.58 | LETM1 | Leucine zipper-EF-hand containing transmembrane protein 1 | 33.35 |
| TRAP1 | TNF Receptor Associated Protein 1 | 1730.73 | | | | PCCB | Propionyl-CoA carboxylase beta chain | 31.27 |
| ACAA2 | Acetyl-CoA acyltransferase 2 | 1220.09 | | | | CEACAM5 | Carcinoembryonic antigen-related cell adhesion molecule 5 | 24.58 |
| AK3 | Adenylate kinase 3 | 981.946 | | | | PTPRF | Protein tyrosine phosphatase, receptor type F | 21.94 |
| UQCRC1 | Ubiquinol-cytochrome c reductase core protein I | 854.604 | | | | MST1R | Macrophage stimulating 1 receptor | 21.13 |
| DCXR | Dicarbonyl/L-xylulose reductase | 819.989 | | | | CNNM3 | Cyclin and CBS domain divalent metal cation transport mediator 3 | 16.02 |
| SSBP1 | Single-stranded DNA-binding protein 1 | 808.072 | | | | SLC27A4 | Fatty acid transporter 4 | 14.53 |

| | | | | | | | | |
|---|---|---|---|---|---|---|---|---|
| NDUFB6 | NADH dehydrogenase [ubiquinone] 1 beta subcomplex 6 | 757.227 | | | | CD70 | CD70 molecule | 14.45 |
| S100A14 | S100 calcium-binding protein A14 | 725.033 | | | | ST14 | Suppression of tumorigenicity 14 (serine peptidase) | 13.58 |
| S100A9 | S100 calcium-binding protein A9 | 680.512 | | | | ECHS1 | Enoyl-CoA hydratase, short chain 1 | 13.28 |
| S100A8 | S100 calcium-binding protein A8 | 621.973 | | | | PVRL4 | Poliovirus receptor-related protein 4 | 13.12 |
| CYB5R1 | Cytochrome b5 reductase 1 | 583.052 | | | | PTPRK | Protein tyrosine phosphatase, receptor type K | 12.76 |
| EPHX1 | Epoxide hydrolase 1 | 581.606 | | | | SPINT1 | Serine peptidase inhibitor, Kunitz type 1 | 12.69 |
| NDUFV1 | NADH dehydrogenase [ubiquinone] flavoprotein 1 | 558.186 | | | | ROBO1 | Roundabout Guidance Receptor 1 | 12.38 |
| COX5A | Cytochrome c oxidase subunit 5A | 558.156 | | | | NIPSNAP1 | Nipsnap homolog 1 | 9.31 |
| SOD2 | Superoxide dismutase 2 | 458.521 | | | | ITGA5 | Integrin subunit alpha 5 | 9.23 |
| GRHPR | Glyoxylate reductase/hydroxypyruvate reductase | 440.284 | | | | ITGAV | Integrin subunit alpha V | 8.62 |
| SDHA | Succinate dehydrogenase complex, subunit A | 393.879 | | | | GOT2 | Glutamate oxaloacetate transaminase 2 | 8.54 |
| CS | Citrate synthase | 381.161 | | | | ALCAM | Activated leukocyte cell adhesion molecule | 7.74 |
| HSD17B10 | Hydroxysteroid (17-beta) dehydrogenase 10 | 369.796 | | | | TUBB2B,TUBB2A | Tubulin beta 2B class IIb; Tubulin beta 2A class IIa | 7.7 |

| | | | | | | | | |
|---|---|---|---|---|---|---|---|---|
| ETFB | Electron transfer flavoprotein subunit beta | 332.828 | | | | PSAP | Prosaposin | 7.51 |
| NDUFA10 | NADH dehydrogenase [ubiquinone] 1 alpha subcomplex 10 | 312.164 | | | | SLC27A1 | Fatty acid transporter 1 | 7.22 |

| **Gene MCF10A cells** | **Protein Name (MCF10A)** | **OC/ND ratio (MCF10A)** | **Gene MCF10AT1 cells** | **Protein Name (MCF10AT1)** | **OC/ND ratio (MCF10AT1)** | **Gene MCF10CA1a cells** | **Protein Name (MCF10CA1a)** | **OC/ND ratio (MCF10CA1a)** |
|---|---|---|---|---|---|---|---|---|
| SCEL | Scellicin | 5318.64 | HTRA2 | High temperature requirement A2 | 7.36 | ACAA2 | Acetyl-CoA acyltransferase 2 | 82.61 |
| ALDH7A1 | Aldehyde dehydrogenase 7 family member A1 | 3111.33 | PMPCB | Peptidase (mitochondrial processing) beta | 5.86 | **ATP2B4** | Plasma membrane calcium-transporting ATPase 4 | 37.51 |
| ECH1 | Enoyl CoA hydratase 1 | 2711.84 | CLPP | ATP-dependent Clp protease | 5.58 | LETM1 | Leucine zipper-EF-hand containing | 33.35 |

| | | | | | | | | |
|---|---|---|---|---|---|---|---|---|
| | | | | proteolytic subunit | | | transmembrane protein 1 | |
| TRAP1 | TNF Receptor Associated Protein 1 | 1730.73 | | | | PCCB | Propionyl-CoA carboxylase beta chain | 31.27 |
| ACAA2 | Acetyl-CoA acyltransferase 2 | 1220.09 | | | | CEACAM5 | Carcinoembryonic antigen-related cell adhesion molecule 5 | 24.58 |
| AK3 | Adenylate kinase 3 | 981.946 | | | | PTPRF | Protein tyrosine phosphatase, receptor type F | 21.94 |
| UQCRC1 | Ubiquinol-cytochrome c reductase core protein I | 854.604 | | | | MST1R | Macrophage stimulating 1 receptor | 21.13 |
| DCXR | Dicarbonyl/L-xylulose reductase | 819.989 | | | | CNNM3 | Cyclin and CBS domain divalent | 16.02 |

| | | | | | | | | |
|---|---|---|---|---|---|---|---|---|
| | | | | | | | metal cation transport mediator 3 | |
| SSBP 1 | Single-stranded DNA-binding protein 1 | 808.0 72 | | | | SLC27A4 | Fatty acid transporter 4 | 14.53 |
| NDUF B6 | NADH dehydrogenase [ubiquinone] 1 beta subcomplex 6 | 757.2 27 | | | | CD70 | CD70 molecule | 14.45 |
| S100 A14 | S100 calcium-binding protein A14 | 725.0 33 | | | | ST14 | Suppression of tumorigenicit y 14 (serine peptidase) | 13.58 |
| S100 A9 | S100 calcium-binding protein A9 | 680.5 12 | | | | ECHS1 | Enoyl-CoA hydratase, short chain 1 | 13.28 |
| S100 A8 | S100 calcium-binding protein A8 | 621.9 73 | | | | PVRL4 | Poliovirus receptor-related protein 4 | 13.12 |

| | | | | | | | | |
|---|---|---|---|---|---|---|---|---|
| CYB5R1 | Cytochrome b5 reductase 1 | 583.052 | | | | PTPRK | Protein tyrosine phosphatase, receptor type K | 12.76 |
| EPHX1 | Epoxide hydrolase 1 | 581.606 | | | | SPINT1 | Serine peptidase inhibitor, Kunitz type 1 | 12.69 |
| NDUFV1 | NADH dehydrogenase [ubiquinone] flavoprotein 1 | 558.186 | | | | ROBO1 | Roundabout Guidance Receptor 1 | 12.38 |
| COX5A | Cytochrome c oxidase subunit 5A | 558.156 | | | | NIPSNAP1 | Nipsnap homolog 1 | 9.31 |
| SOD2 | Superoxide dismutase 2 | 458.521 | | | | ITGA5 | Integrin subunit alpha 5 | 9.23 |
| GRHPR | Glyoxylate reductase/hydroxypyruvate reductase | 440.284 | | | | ITGAV | Integrin subunit alpha V | 8.62 |

| | | | | | | | | |
|---|---|---|---|---|---|---|---|---|
| SDHA | Succinate dehydrogenase complex, subunit A | 393.879 | | | | GOT2 | Glutamate oxaloacetate transaminase 2 | 8.54 |
| CS | Citrate synthase | 381.161 | | | | ALCAM | Activated leukocyte cell adhesion molecule | 7.74 |
| HSD17B10 | Hydroxysteroid (17-beta) dehydrogenase 10 | 369.796 | | | | TUBB2B, TUBB2A | Tubulin beta 2B class IIb; Tubulin beta 2A class IIa | 7.7 |
| ETFB | Electron transfer flavoprotein subunit beta | 332.828 | | | | PSAP | Prosaposin | 7.51 |
| NDUFA10 | NADH dehydrogenase [ubiquinone] 1 alpha subcomplex 10 | 312.164 | | | | SLC27A1 | Fatty acid transporter 1 | 7.22 |

**Supplementary Table 2.** Top 25 genes and corresponding proteins that exhibited more than a 100-fold increase in plasma membrane association under OC conditions compared to ND conditions in MCF10A cells (left), and more than a 5-fold increase in MCF10AT1 (middle) and MCF10CA1a (right) cells, as identified by mass spectrometry. One ion transporter showing plasma membrane relocation under OC conditions in MCF10CA1a cells is highlighted in bold.

.

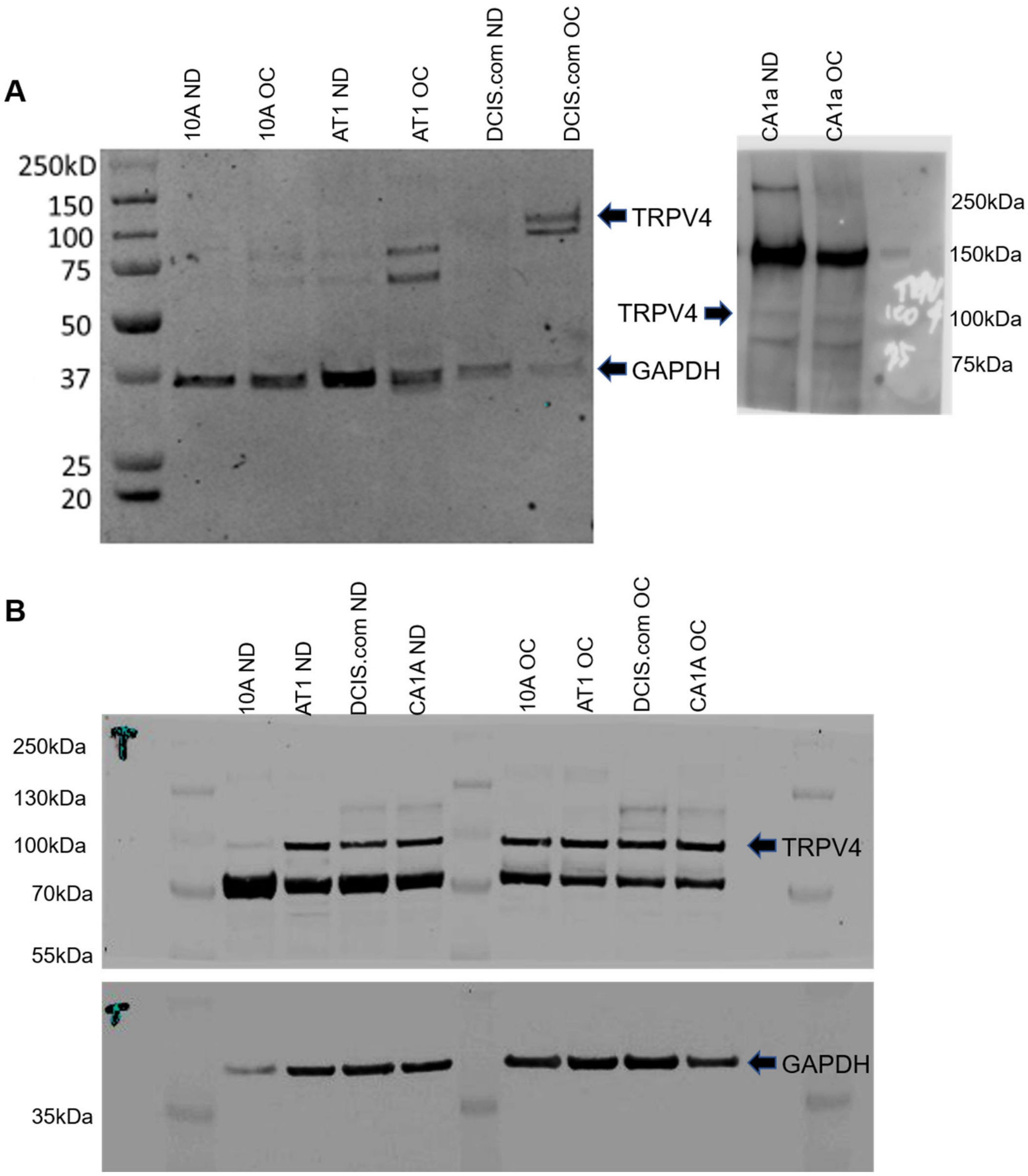


**Supplementary Figure 5. Original immunoprecipitation and western blot images. (A)** Left: Plasma membrane proteins pulled down after cell surface biotinylating with streptavidin beads and immunoblotted for TRPV4 and GAPDH (loading control) for ND vs OC cells of MCF10A (10A), MCF10AT1 (AT1), and MCF10DCIS.com (DCIS.com). Right: The same procedure was performed to compare PM TRPV4 between ND vs OC MCF10CA1a cells. (**B)** Overall TRPV4 protein levels from whole-cell lysates from four 10A cell derivatives. GAPDH was a loading control.

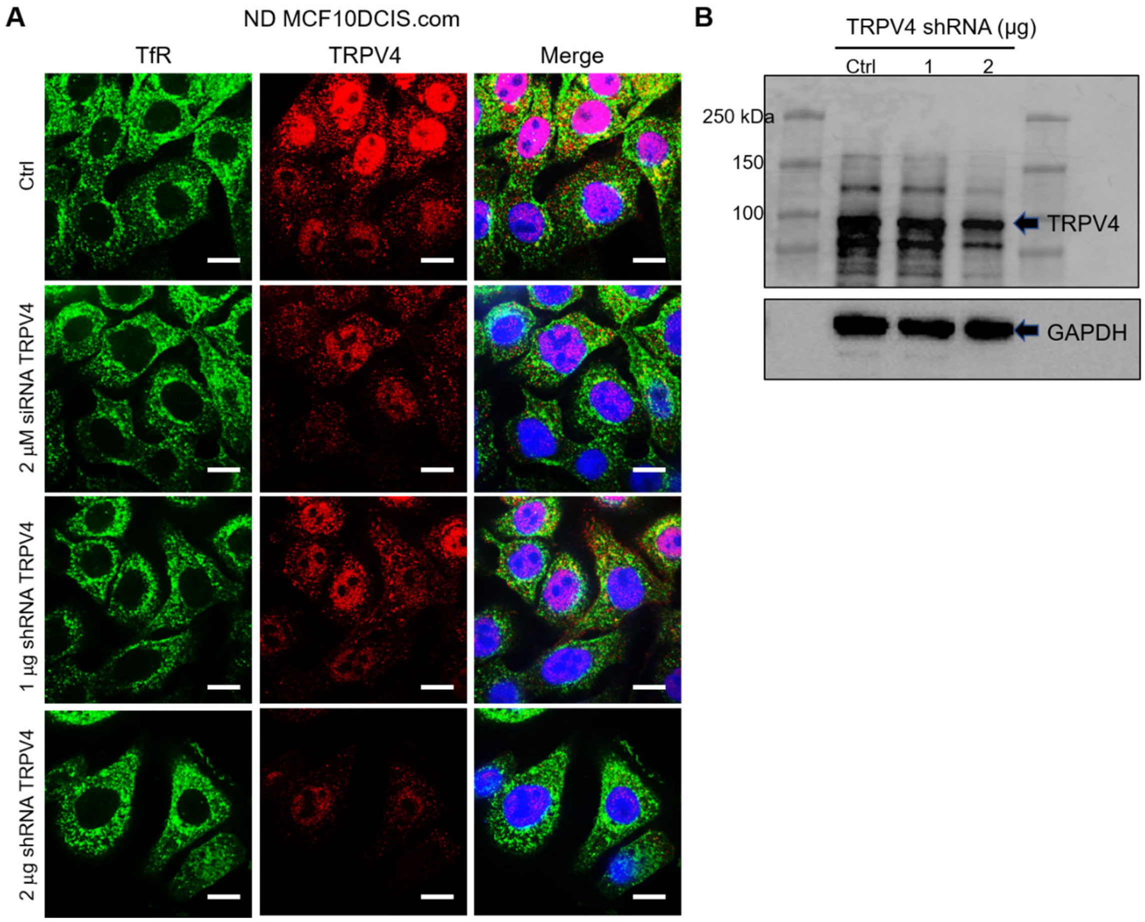


**Supplementary Figure 6. Binding specificity of TRPV4 antibody.** Immunofluorescence images (**A**) and immunoblots (**B**) verified the binding specificity of TRPV4 antibodies in control ND DCIS.com cells and TRPV4-depleted cells treated with either 2 µM siRNA (Dharmacon On Target Plus SMART Pool L-004195-00-0005) or 1 and 2 µg shRNA (shRNA pool, Santa Cruz sc-61726-SH) for 36 hr. (**A**) Compared to invariant transferrin receptor (TfR) staining (green), TRPV4 (red) depletion was 40% with 1 µg shRNA and 80% with 2 µg shRNA, as quantified by intensity measurements. DAPI (blue) is also shown in the merged images. All images were visualized using the same intensity settings. Scale bar = 20 µm. (**B**) Immunoblot results confirmed this dose-responsive depletion of TRPV4, with 33% reduction observed at 1 µg shRNA and 51% at 2 µg.

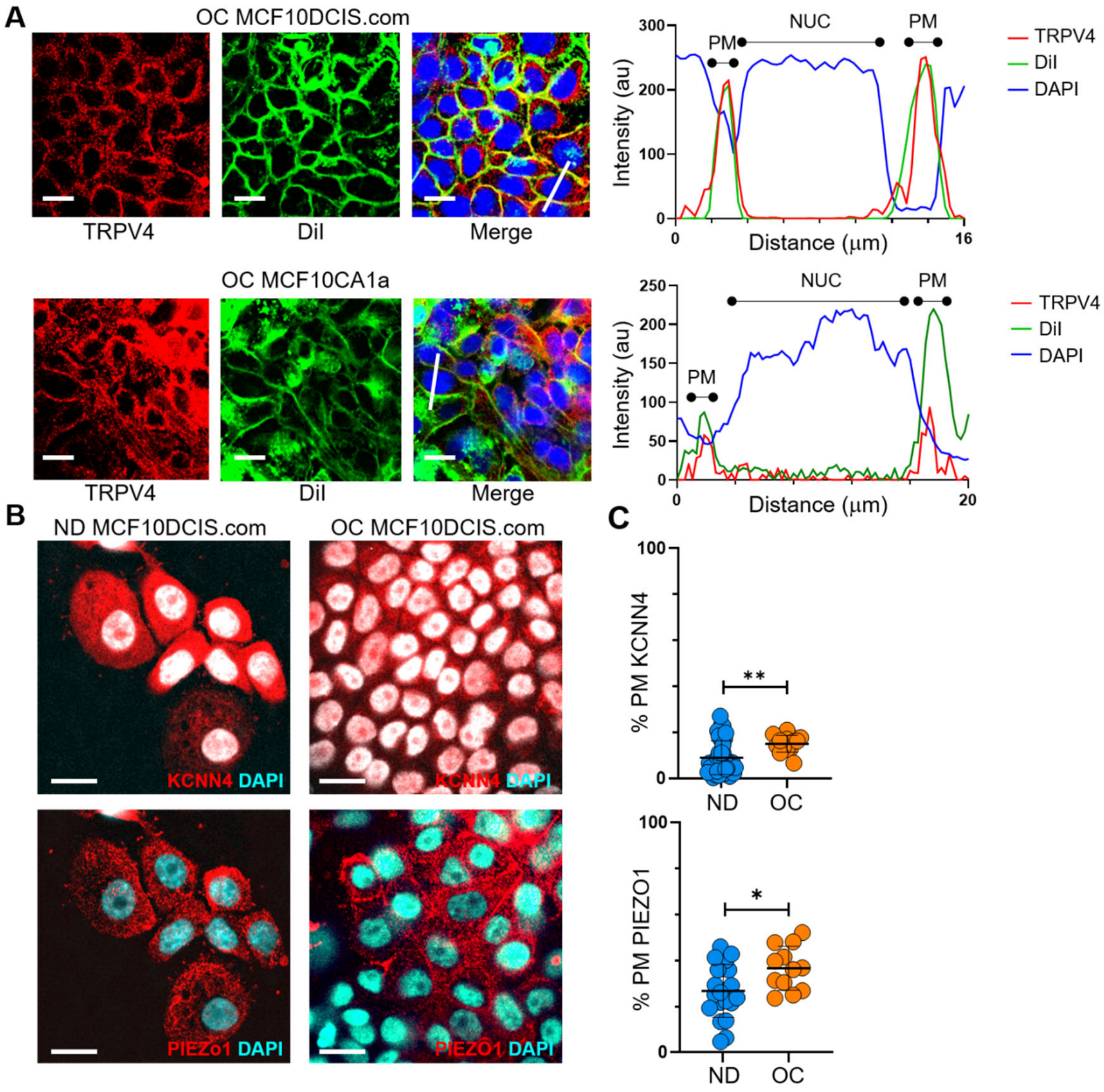


**Supplementary Figure 7.** Various ion channels are relocated to the plasma membrane under cell crowding conditions. (**A**) We used the plasma membrane marker DiIC18(3) to confirm the association of TRPV4 with the plasma membrane in MCF10DCIS.com and MCF10CA1a cells under OC conditions. As described in the Methods section, we stained DiIC18(3) in live cells and co-stained TRPV4 and DAPI in fixed and permeabilized cells. The IF images show TRPV4 (red), DiIC18(3) (DiI, green), and DAPI (blue). The line profile plots on the right demonstrate colocalization of TRPV4 with DiIC18(3) at the plasma membrane (PM), marked by the green DiIC18(3) signal, which overlaps with the red TRPV4 signal. The nucleus location (NUC) is indicated by the blue DAPI signal. Scale bar = 20 μm. (**B**) We examined the relocation of KCNN4 and PIEZO1 to the plasma membrane in response to cell crowding. Mass spectrometry showed a slight increase in KCNN4 at the plasma membrane under OC conditions. In ND MCF10DCIS.com cells, KCNN4 was predominantly cytosolic, whereas PIEZO1 showed some plasma membrane association. Under OC conditions, both KCNN4 and PIEZO1 showed a modest relocation to the plasma membrane. (**C**) Line analysis confirmed a slight increase in plasma membrane association for both KCNN4 and PIEZO1 under OC conditions compared to ND conditions. Scale bar = 20 μm. For the statistical analysis, we employed a nonparametric approach using the Mann-Whitney test with a two-tailed p-value. The levels of statistical significance are denoted as follows: **** indicates $p < 0.0001$, *** indicates $p < 0.001$, * indicates $p < 0.1$, and "ns" indicates $p > 0.05$.

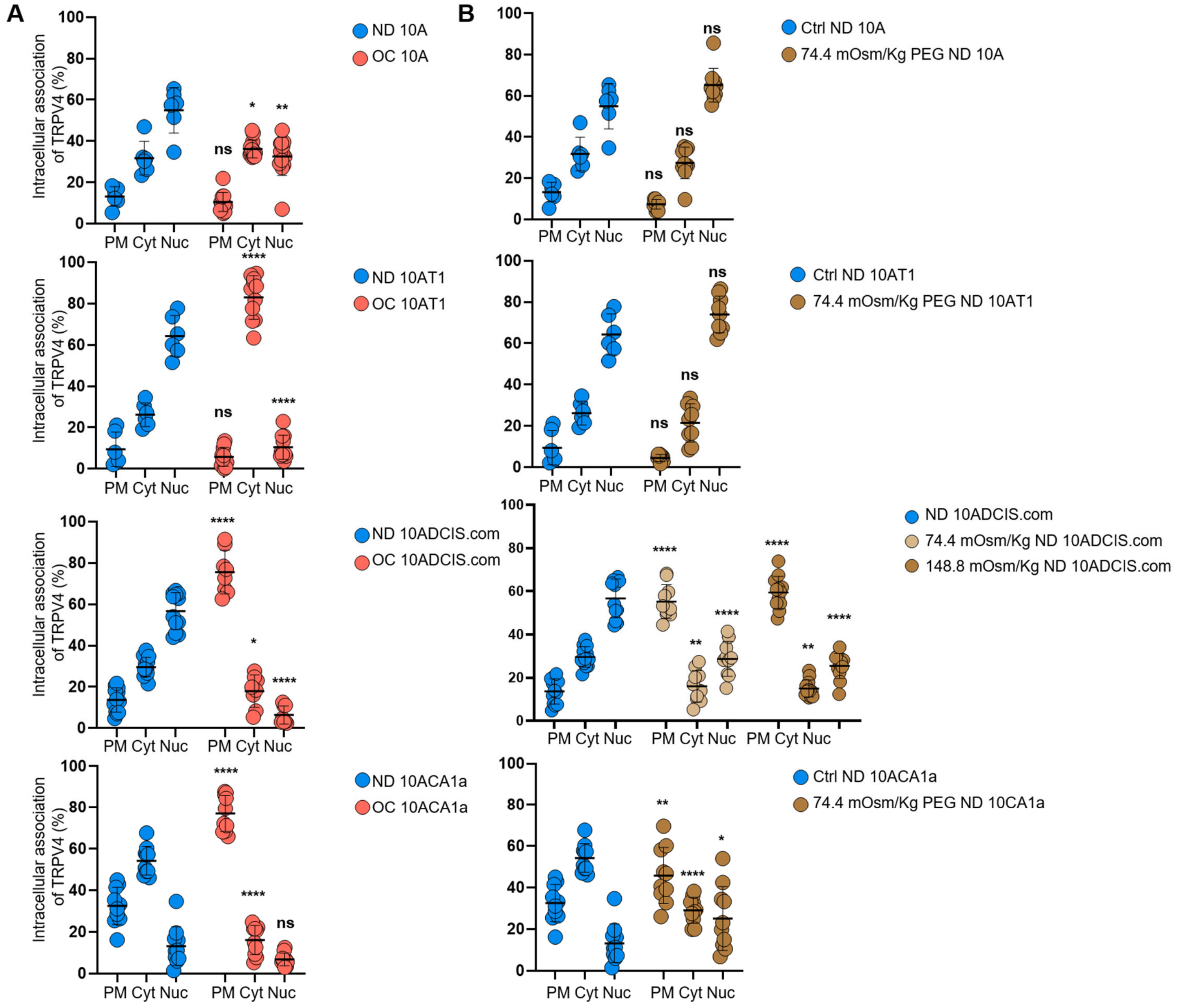


**Supplementary Figure 8. Plots of the relative intracellular TRPV4 associations between ND and OC or hyperosmotic conditions in all four cell types. (A)** Relative TRPV4 associations with the plasma membrane (PM), cytoplasm (Cyt), and nucleus (Nuc) are plotted for ND versus OC conditions in MCF10A (10A), MCF10AT1 (10AT1), MCF10DCIS.com (10DCIS.com), and MCF10CA1a (10CA1a) cells. **(B)** Similar analyses were performed to compare the intracellular TRPV4 associations in PM, Cyt, and Nuc between control ND and 74.4 or 148.8 mOsm/kg PEG

300 treatment groups. We employed a nonparametric approach using the Mann-Whitney test with a two-tailed p-value for the statistical analysis. The levels of statistical significance are denoted as follows: **** indicates $p < 0.0001$, *** indicates $p < 0.001$, * indicates $p < 0.1$, and "ns" indicates $p > 0.05$.

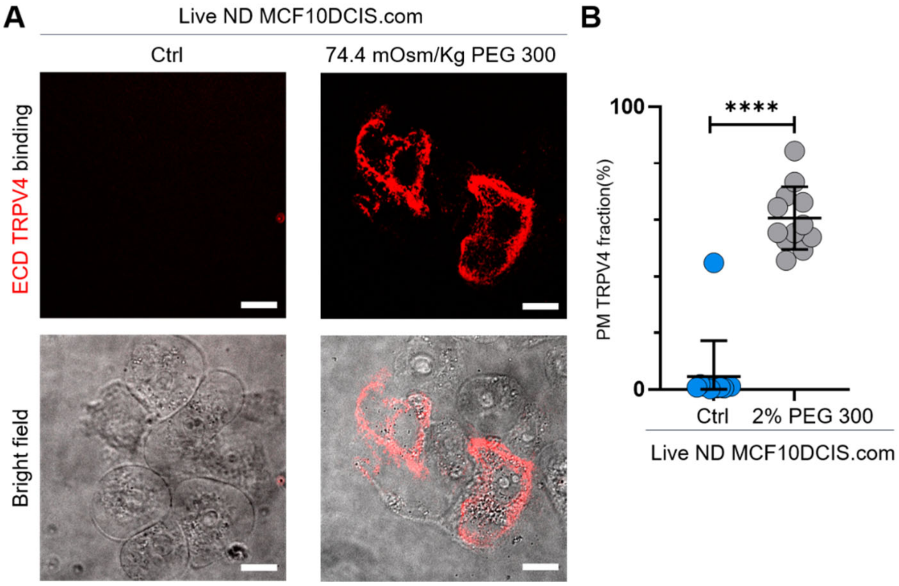


**Supplementary Figure 9. Validation of mechanosensitive relocation of TRPV4 to the plasma membrane in MCF10DCIS.com cells. (A)** To determine whether the seemingly plasma membrane-associated TRPV4 under hyperosmotic conditions was indeed localized at the plasma membrane of live cells, we used an extracellular domain (ECD) TRPV4 antibody. See the Methods section for sample preparation details. Live-cell imaging with 60X confocal microscopy revealed distinct binding of the ECD TRPV4 antibodies (red) exclusively in MCF10DCIS.com cells treated with 74.4 mOsm/kg PEG 300 for 15 minutes. No binding was observed in the untreated control group. Brightfield images are shown below the fluorescence images. **(B)** Line analysis demonstrated that ~60% of TRPV4 was plasma membrane-associated under 2% (74.4 mOsm/kg) PEG 300 conditions, compared to ~0% in untreated cells. Statistical analysis was conducted using a nonparametric Mann-Whitney test with two-tailed p-values. Statistical significance levels are indicated as follows: **** $p < 0.0001$, *** $p < 0.001$, * $p < 0.1$, and "ns" denotes $p > 0.05$.

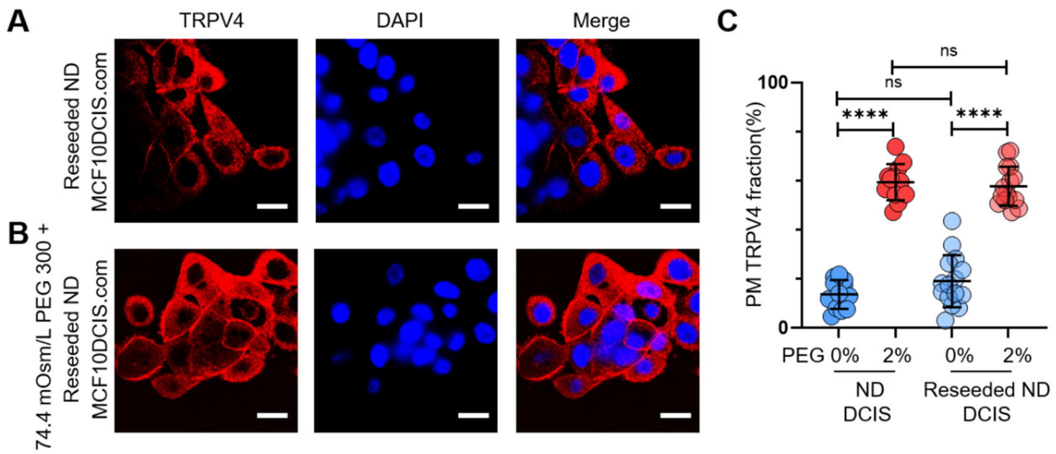


**Supplementary Figure 10. Reversibility of cell-crowding-driven mechanosensing effects in MCF10DCIS.com cells. (A)** Previously OC MCF10DCIS.com cells were trypsinized and reseeded under ND conditions. These cells were fixed and stained for TRPV4 (red) and DAPI (blue) for IF imaging. While TRPV4 was elevated at the plasma membrane under OC conditions, it redistributed to the cytoplasm after the cells were reseeded under ND conditions. **(B)** The reseeded ND cells responded similarly to hyperosmotic stress (74.4 mOsm/Kg PEG 300 for 15 min), with TRPV4 relocating to the plasma membrane. **(C)** Line analysis demonstrated that the reseeded ND cells exhibited a TRPV4 distribution pattern comparable to their initial ND counterparts without (blue circles) and with (red circles) hyperosmotic conditions. Statistical analysis was performed using a nonparametric Mann-Whitney test with two-tailed p-values. Statistical significance levels are denoted as follows: **** $p < 0.0001$, *** $p < 0.001$, * $p < 0.1$, and "ns" indicates $p > 0.05$.

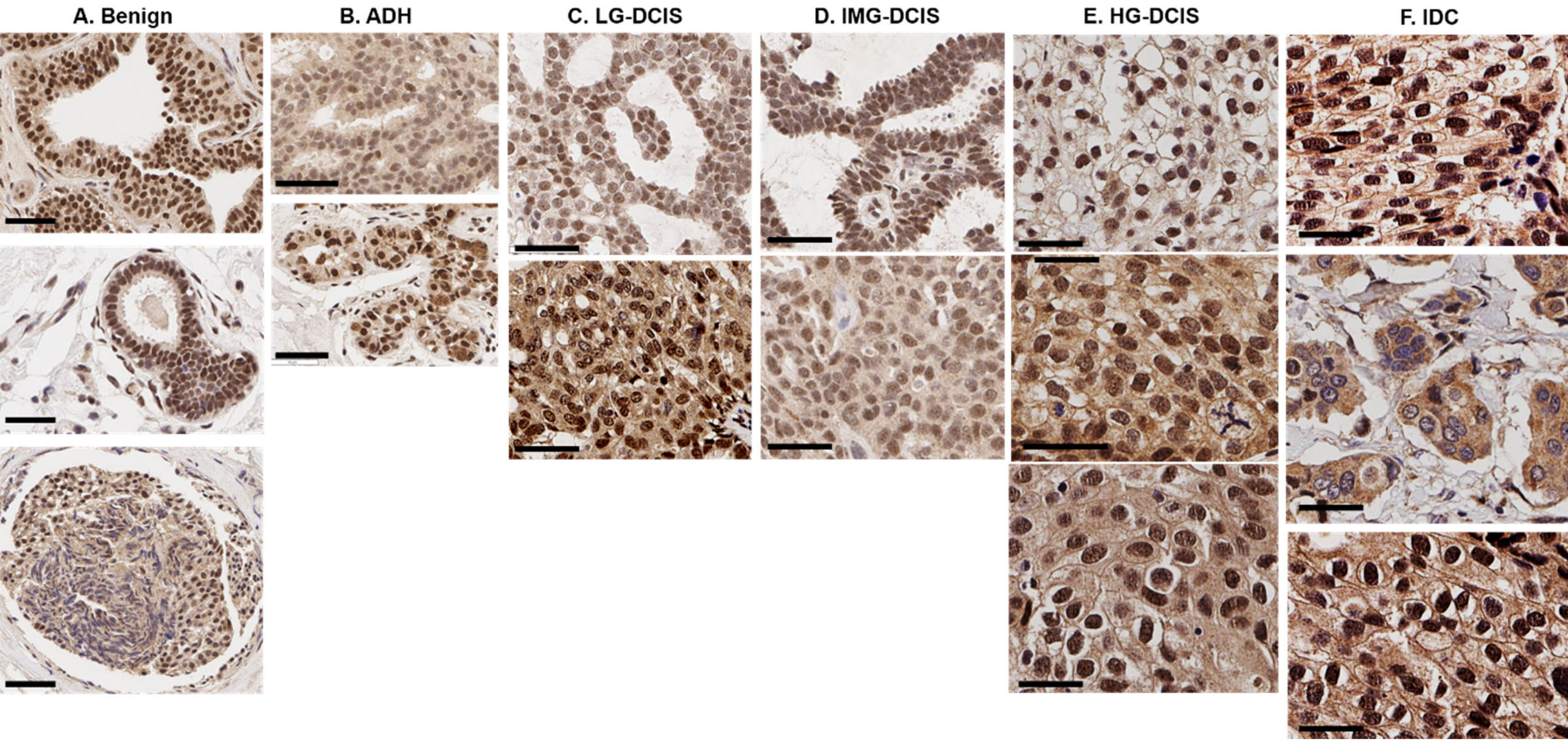


**Supplementary Figure 11. Pathology-dependent differential TRPV4 distributions in patients' IHC images.** Representative IHC images for each pathology from different patients are displayed. A selective presence of TRPV4 pools in the plasma membrane was mainly observed in high-grade DCIS and IDC lesions. Two pathologists independently conducted annotations. When a consensus was not reached, the case was labeled "equivocal." The cases were categorized according to the subsequent criteria:

Case 1: Absence of TRPV4.

Case 2: Intracellular TRPV4 localization.

Case 3: Presence of TRPV4 in the plasma membrane, with or without intracellular TRPV4.

**A. Benign cases.**

IHC images - <u>(Top)</u>: Pathology: UDH: usual ductal hyperplasia; protein distribution: case 2;

<u>(Middle)</u>: Pathology: benign (columna); protein distribution: case 2;

(Bottom): Pathology: benign (papilloma); protein distribution: case 2.

Scale bars = 30 μm.

**B. Atypical ductal hyperplasia (ADH) cases.**

IHC images - (Top): Pathology: ADH: atypical ductal hyperplasia; protein distribution: case 2;

(Bottom): Pathology: ADH: atypical ductal hyperplasia; protein distribution: case 2.

Scale bars = 50 μm.

**C. Low-grade (LG) DCIS cases.**

IHC images - (Top): Pathology: low-grade DCIS; protein distribution: case 2;

(Bottom): Pathology: low-grade DCIS; protein distribution: case 2.

Scale bars = 50 μm.

**D. Intermediate-grade (IMG) DCIS cases.**

IHC images - (Top): Pathology: intermediate-grade DCIS; protein distribution: case 2;

(Bottom): Pathology: intermediate-grade DCIS; protein distribution: case 2.

Scale bars = 50 μm.

**E. High-grade (HG) DCIS cases.**

IHC images - (Top): Pathology: high-grade DCIS; protein distribution: case 3;

(Middle): Pathology: high-grade DCIS; protein distribution: the distinction between case 2 and case 3 is equivocal;

(Bottom): Pathology: high-grade DCIS; protein distribution: case 3.

Scale bars = 50 μm.

**F. Invasive ductal carcinoma (IDC) cases.**

IHC images - (Top): Pathology: IDC; protein distribution: case 3;

(Middle): Pathology: IDC; protein distribution: case 2.

(Bottom): Pathology: IDC; protein distribution: case 3.

Scale bars = 50 μm.

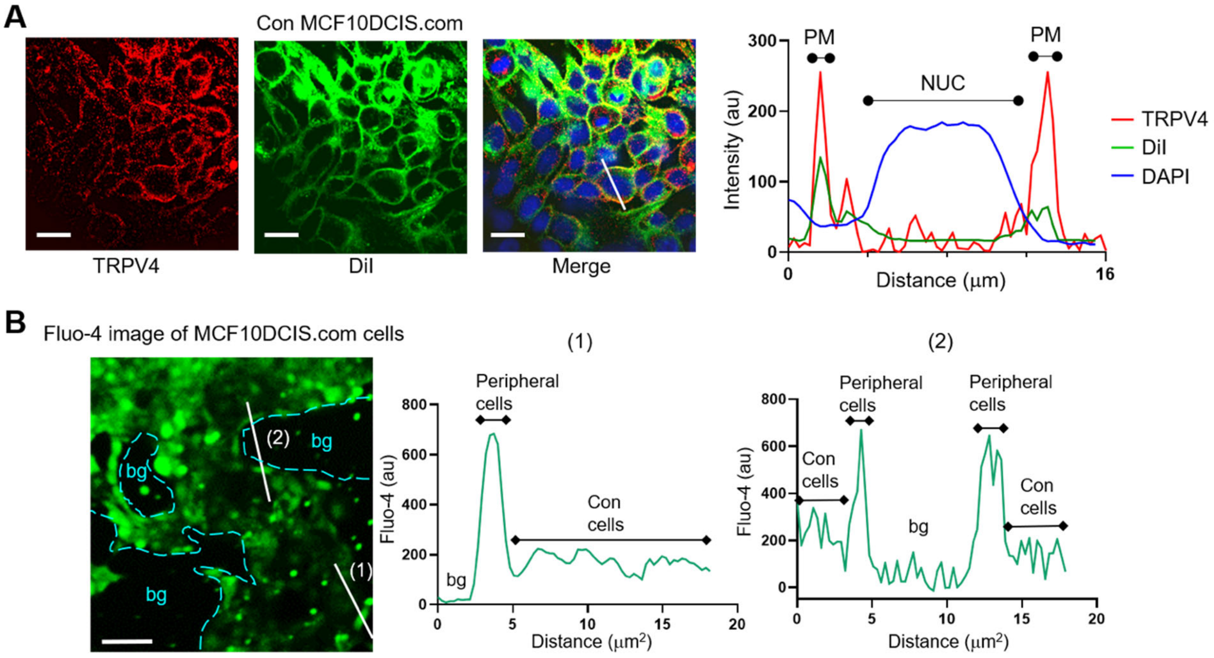


**Supplementary Figure 12. Peripheral cells within MCF10DCIS.com cell clusters exhibit higher calcium levels due to reduced cell crowding effects. (A)** Confluent cell density (Con) also triggered the relocation of TRPV4 (red) to the plasma membrane, similar to OC conditions, as shown in IF image (left). We used DiIC18(3) to indicate the plasma membrane location (green; DiI; middle image), and the merged image (right) of TRPV4 and DiIC18(3) shows excellent overlay at the plasma membrane (PM), as illustrated in the line profile plots from our line analysis. DAPI (blue) staining was used to locate the nucleus (NUC). Scale bar = 20 μm. **(B)** Confluent cell density resulted in lower intracellular calcium levels compared to less confluent cells. Live MCF10DCIS.com cells stained with Fluo-4 were imaged using confocal microscopy at 488 nm. Two line profiles (1, 2) crossing peripheral cells (less confluent than confluent cells) and adjacent confluent cells within the clusters clearly showed that the peripheral cells have a higher Fluo-4 signal (700 au) compared to the confluent cells within the cluster (200 au), highlighting the crowding-induced intracellular calcium reduction. Background regions (bg) were noted with cyan dashed-lines in the fluorescent image and in the plots. Scale bar = 100 μm.

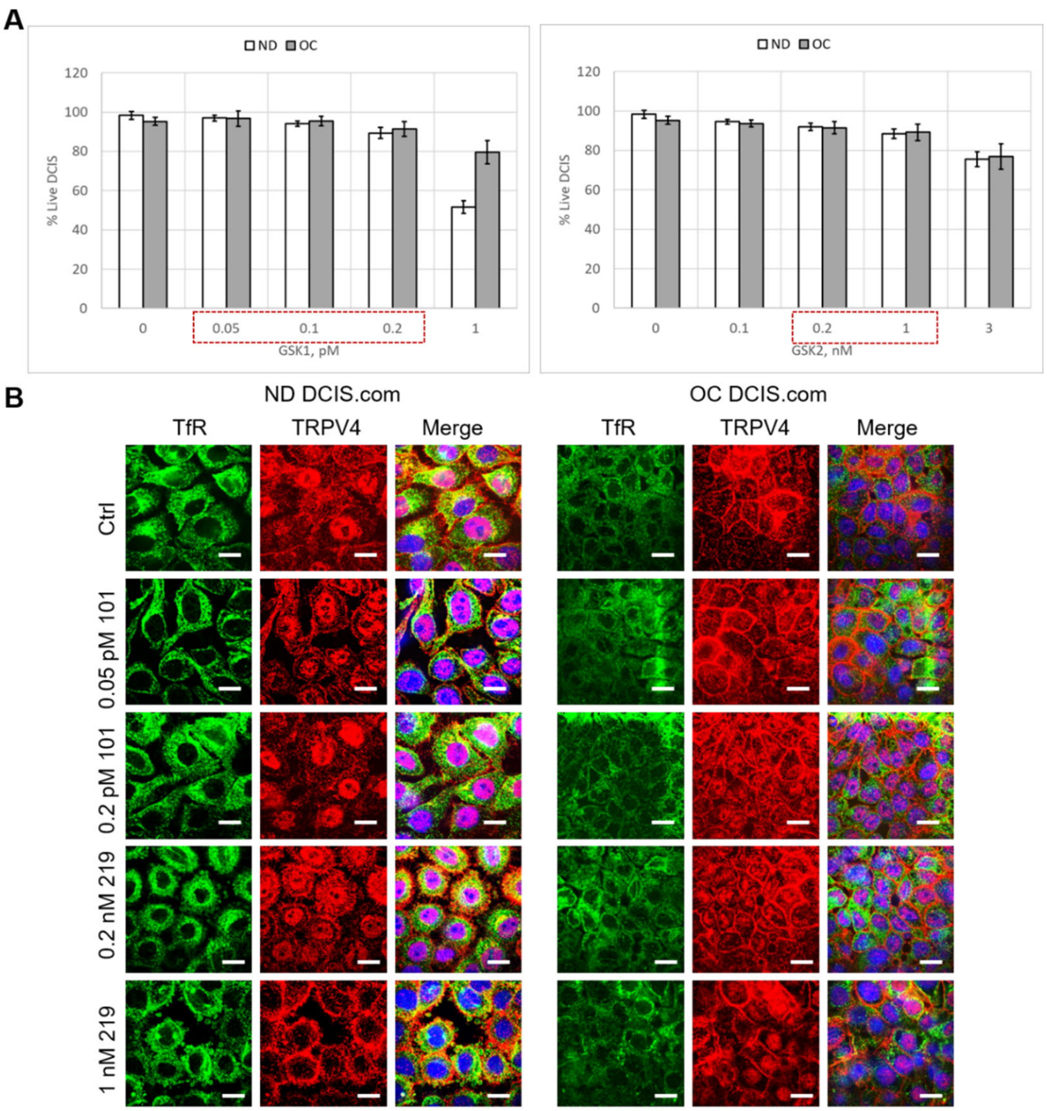


**Supplementary Figure 13. Determination of treatment concentration ranges for TRPV4 activator (GSK101) and inhibitor (GSK219). (A)** Cell viability assays in which viable cells were counted based on trypan blue staining after two days of GSK101 or GSK219 treatment in the specified concentration ranges of DCIS in ND (white bars) and OC (gray bars) conditions. Treatment ranges were selected so that cell viability was >90%. Concentrations used for dose-dependent assays were 0.05 and 0.2 pM for GSK101, and 0.2 and 1 nM for GSK219 (marked by dotted red boxes). **(B)** Representative confocal microscopy immunofluorescence images showed effects of GSK101 (0.05 and 0.2 pM) or GSK219 (0.2 and 1 nM) treatment for two days on TRPV4 (red) and control transferrin receptor (TfR; green) distributions in ND and OC cells in a dose-dependent manner. DAPI (blue) signal is shown in the merged images. Scale bar = 20 μm.

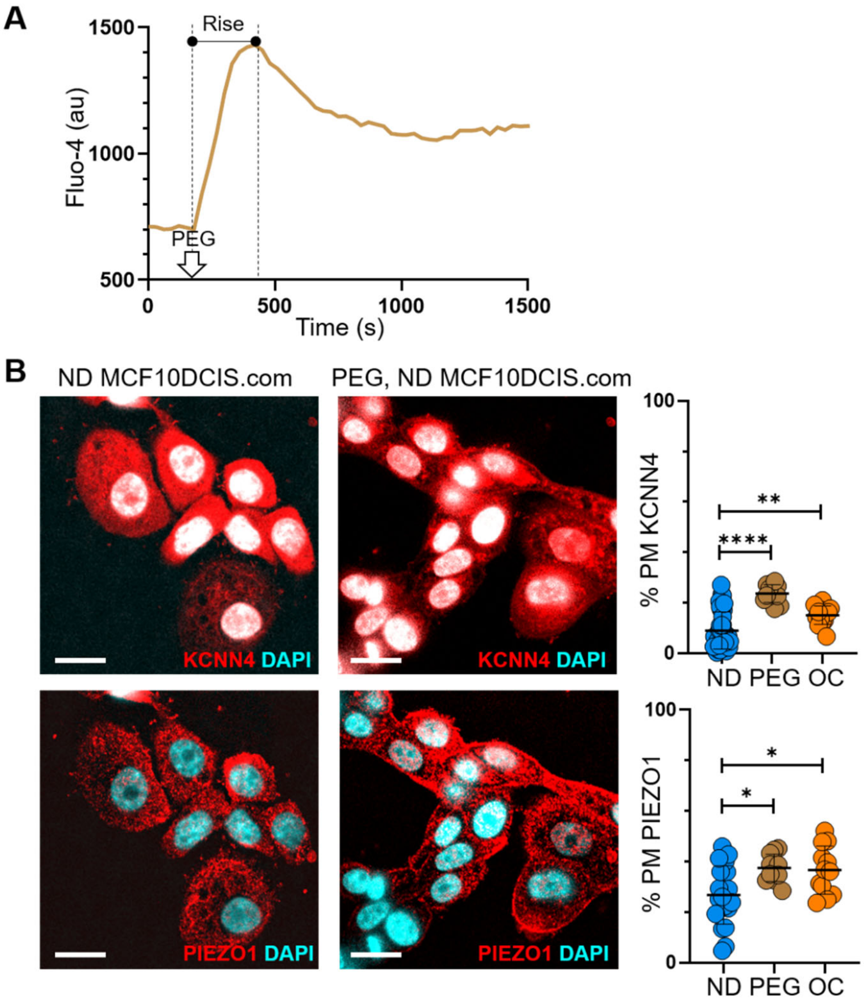


**Supplementary Figure 14. Hyperosmotic stress also induces plasma membrane relocation of ion channels, similar to cell crowding. (A)** Using the Fluo-4 assay, we observed an initial calcium spike (marked as "Rise") in ND MCF10DCIS.com cells in response to 74.4 mOsmol/kg PEG 300, due to osmotic water outflow. This was followed by a homeostatic relaxation, aimed at restoring calcium levels, which likely involved the inhibition of ion channels like TRPV4, leading to their plasma membrane relocation. Scale bars = 20 μm. **(B)** The same hyperosmotic condition (74.4 mOsm/Kg PEG 300 for 15 min) led to the relocation of KCNN4 and PIEZO1 to the plasma membrane, similar to the relocations observed under OC conditions. Line analysis results showed the relative relocations of each channel in response to hyperosmotic (PEG) and cell crowding (OC) stresses, compared to ND conditions.

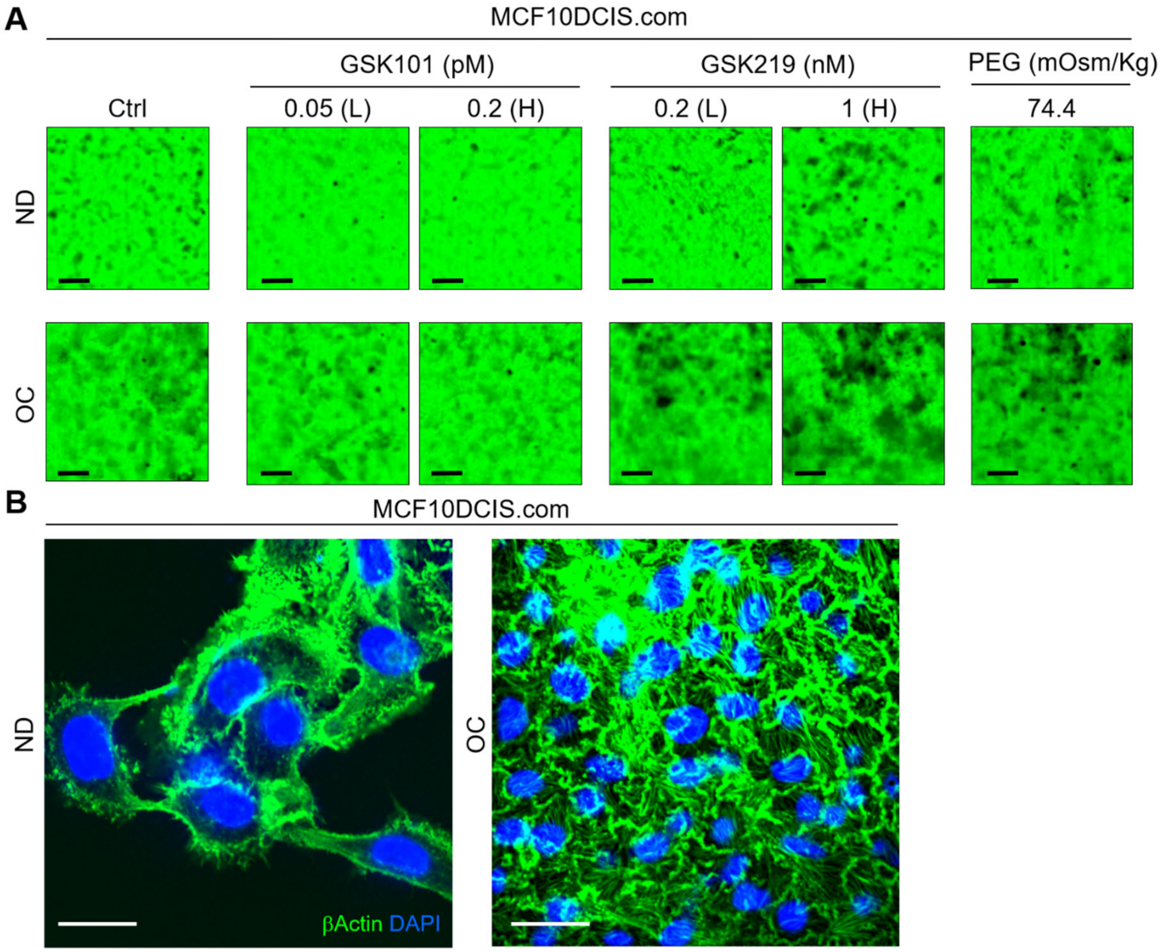


**Supplementary Figure 15. Mechanical stresses and TRPV4 activation status affect MCF10DCIS.com cell invasiveness. (A)** The effects of 2 days of treatment with GSK101 (0.05 and 0.2 pM), GSK219 (0.2 and 1 nM), and PEG 300 (74.4 mOsm/kg) on the invasiveness of MCF10DCIS.com cells under ND and OC conditions. Gelatin488 images demonstrated the dose-dependent negative effects of GSK101 and positive effects of GSK219 on cell invasiveness. Similar to GSK219, PEG 300 also increased cell invasiveness. Scale bar = 100 µm. **(B)** IF images of β–actin (green) and DAPI (blue) in MCF10DCIS.com cells under ND (left) and OC (right) conditions. Strong stress fiber formation was observed in OC cells, while it was absent in ND cells, which was reflected by the increased stiffness of OC cells (**Fig. 2C**). This suggests that cell crowding may enhance cell motility. Scale bar = 20 µm.

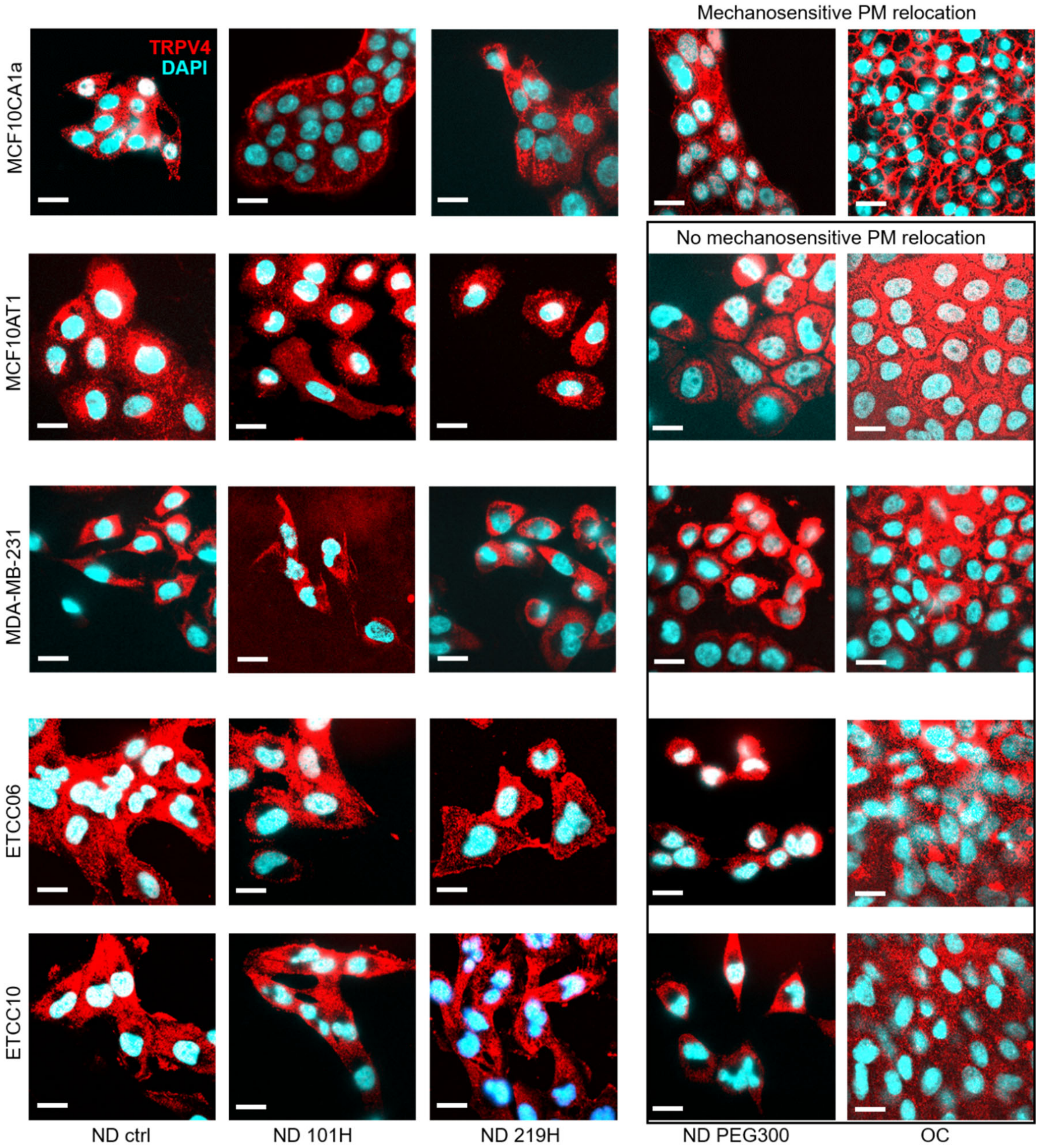


**Supplementary Figure 16. Cells lacking the capability for activating pro-invasive mechanotransduction pathway via TRPV4 inhibition-induced cell volume reduction do not relocate TRPV4 to the plasma membrane under TRPV4 inhibition and mechanical stresses.** The effects of treatment with GSK101 (0.2 pM, 1-hour), GSK219 (1 nM, 1-hour), and PEG 300 (74.4 mOsm/kg, 15-miniutes) on cells under ND or OC conditions revealed differences in TRPV4 localization (red) in the immunofluorescence (IF) images (cyan: DAPI). Only MCF10CA1a cells show GSK219, PEG 300, and OC-induced TRPV4 relocation to the plasma membrane. Other cell types, including MCF10AT1, MDA-MB-231, ETCC-06, and ETCC-10, did not exhibit this translocation. Scale bar = 20 μm.
